# Development of a continuous fluorescence-based assay for N-terminal acetyltransferase D

**DOI:** 10.1101/2020.12.24.424328

**Authors:** Yi-Hsun Ho, Lan Chen, Rong Huang

**Author notes:** Corresponding author: Rong Huang.

## Abstract

N-terminal acetylation catalyzed by N-terminal acetyltransferases (NATs) has various biological functions in protein regulation. N-terminal acetyltransferase D (NatD) is one of the most specific NAT with only histone H4 and H2A proteins as the known substrates. Dysregulation of NatD has been implicated in colorectal and lung cancer progression, implying its therapeutic potential in cancers. However, there is no reported inhibitor for NatD yet. To facilitate the discovery of small-molecule NatD inhibitors, we report the development of a fluorescence-based acetyltransferase assay in 384-well high-throughput screening (HTS) format through monitoring the formation of coenzyme A. The fluorescent signal is generated from the adduct in the reaction between coenzyme A and fluorescent probe ThioGlo4. The assay exhibited a Z’-factor of 0.77 and a coefficient of variation of 6%, indicating it is a robust assay for HTS. A pilot screen of 1280 pharmacologically active compounds and subsequent validation identified two hits, confirming the application of this fluorescence assay in HTS.

## Introduction

N-terminal acetylation has been detected on 80–90% of human cytosolic and transmembrane proteins,^1,2^ which impacts diverse biological processes, such as regulating protein degradation, protein translocation, and protein-protein interactions.^3^ This modification is catalyzed by N-terminal acetyltransferase (NATs), which transfer the acetyl group from acetyl-coenzyme A (AcCoA) to the alpha-N-terminal amino group of the protein substrates.^4^ In humans, seven different NATs (NatA–NatF, NatH) have been identified concerning their subunit compositions and substrate specificity profiles.^5^ Among those seven NATs, substrate profiling revealed that NatD is a unique NAT enzyme because of its substrate-specific for histones H4 and H2A that share an identical SGRGK sequence at the N-terminus.^6–9^ Although the function of N-terminal acetylation on H4 and H2A remains obscure, NatD has been implicated in cancer cell growth, migration, and invasion in colorectal and lung cancers.^10–12^ Depletion of NatD in colorectal cancer cells induced the mitochondrial caspase-9-mediated apoptosis without affecting non-cancerous fibroblasts.^10^ Moreover, the loss of NatD sensitized the colorectal cancer xenograft mice to the 5-fluorouracil treatment to slow the tumor growth through transcriptional downregulation of protein arginine methyltransferase 5 (PRMT5).^11^ In lung cancer, NatD expression level is inversely proportional to the survival of lung cancer patients.^12^ NatD has been reported to promote the migration and invasion of lung cancer cells both *in vitro* and *in vivo*, while the knockdown of NatD suppressed Slug’s transcription factor and subsequently repressed the epithelial-to-mesenchymal transition (EMT) in lung cancer cells.^12^ Although the acetyltransferase activity of NatD is implicated in the aforementioned diseases, there is no inhibitor being reported for NatD yet. Therefore, there is a need to discover potent and specific inhibitors for NatD to elucidate its function.

High throughput screening (HTS) has become a popular method to identify initial hit compounds. To date, there are two different types of assays reported to characterize the N-terminal acetylation *in vitro*. First is the radioactive assay using [^14^C]-AcCoA to study the N-terminal acetylation status on peptide substrates through high-performance liquid chromatography (HPLC)-based method or scintillation.^13–15^ However, the low throughput of HPLC and requirement of special handling of radioactive materials limit its application for HTS. To date, there is only radioactive assay has been reported for detecting NatD catalytic activity.^9^ Another assay is to detect the production of CoA using the 5,5′-dithiobis-(2-nitro-benzoic acid) (DTNB). Briefly, the thiol group of CoA reacts with DTNB to form a disulfide and release 5-thio-2-nitrobenzoic acid (TNB), which can be quantified by measuring its absorbance at 412 nm.^16^ This DTNB assay is relatively fast, but it has a CoA detection limit at single-digit micromolar because it requires higher substrate concentrations to achieve at least 10 μM of CoA formation to obtain reasonable standard deviations.^17^ Since the K_m_ values of substrates of NatD are below 10 μM, DTNB assay is not suitable for the detection of CoA formation under NatD catalyzed N-terminal acetylation.

Herein, we report the development of a continuous fluorescence-based assay to quantify CoA production by forming a fluorescent adduct with the thiol fluorescent probe IV (ThioGlo4).^18^ Moreover, we optimized this fluorescence-based assay in a 384-well microplate with a Z’-factor of 0.77 and a coefficient of variation (CV) of 6%, demonstrating its applicability for HTS.

## Materials and Methods

### Materials

Acetyl coenzyme A lithium salt (AcCoA) and coenzyme A sodium salt hydrate (CoA) were purchased from Sigma (St. Louis, MO). ThioGlo4 was purchased from Berry and Associates Inc., Dexter, MI. Nickel-nitrilotriacetic acid (Ni-NTA) resin was purchased from GE Healthcare. The plasmid encoded an N-terminal 8X His-SUMO-tagged hNatD in the pET-21a vector was obtained from Genscript. Library of Pharmaceutically Active Compounds (LOPAC, 1280) from Sigma was housed at Chemical Genomics Facility at Purdue Institute for Drug Discovery.

### Expression and Purification of Recombinant Human NatD

The full-length human NatD gene was amplified and cloned into a modified pET-21a(+) vector containing an 8x-His-tag and SUMO protease cleavage site at the N-terminus (Genscript). This pET-21a(+)-hNatD construct was transformed into *Escherichia coli* BL21 (DE3) codon plus RIL cells in Luria Broth (LB) media containing 50 μg/mL ampicillin, which were grown an OD_600_ of 0.7 at 37°C and induced with 1 mM isopropyl β-D-1-thiogalactopyranoside (IPTG) at 16 °C overnight. Cells were harvested by centrifugation at 4200 rpm for 30 min at 4 °C. The cell pellets were resuspended in lysis buffer containing 20 mM Tris, pH 7.5, 300 mM NaCl, 5 mM β-mercaptoethanol (BME), and sonicated at an amplitude of 40% for 15 min, oscillating between 2-sec sonication and 5-sec break. The lysate was clarified by centrifugation at 12000 rpm at 4 °C for 30 min. The clear lysate was loaded onto the HisTrap FF column and washed with lysis buffer supplemented with 20 mM imidazole. The His-SUMO-tagged hNatD was eluted in the lysis buffer containing 300 mM imidazole. SUMO protease was added to 8XHis-SUMO-tagged hNatD in a weight ratio of 1:50 and gently shaken at 4 °C to remove the 8XHis-SUMO-tag from hNatD. The resultant solution was run over a HisTrap FF column to remove SUMO protease, 8XHis-SUMO-tag, and any uncut His-SUMO-hNatD. Eluted fractions containing untagged hNatD were pooled and dialyzed into a buffer containing 50 mM Tris, pH 7.5, 100 mM NaCl, and 0.5 mM TCEP. After purification through a HiPrep 16/60 Sephacryl S-100 size-exclusion column (SEC), the peak fractions were pooled and concentrated using Amicon Ultra 15 mL concentrator with 10 kDa cut-off (MWCO; Millipore). The concentration was determined by Nanodrop, and purity was confirmed to be greater than 95% pure via SDS-PAGE.

### Synthesis of Peptide Substrates

Peptides representing H4-8 (SGRGKGGK) and H4-21 (SGRGKGGKGLGKGGAKRHRKV) were synthesized on Rink Amide MBHA resin following a standard Fmoc protocol on a CEM Liberty Blue microwave peptide synthesizer. Standard couplings of amino acids were carried out at 0.2 M in DMF and external amino acids at 0.1 M in DMF using 0.5 M DIC and 1.0 M Oxyma in DMF for activation. Fmoc protection groups were removed by 20% (v/v) piperidine in DMF. The resin was transferred to a filter-equipped syringe, washed with CH_2_Cl_2_ (3 mL) and MeOH (3 mL) alternatively three times, and dried under air. The peptides were cleaved from the resin in the presence of a cleavage cocktail (5 mL) containing trifluoroacetic acid (TFA)/2,2’-(Ethylenedioxy)-diethanethiol (DODT) /triisopropylsilane (TIPS) /water (94:2.5:1:2.5 v/v) at room temperature for 4 h. Volatiles of the filtrate was removed under N_2,_ and the residue was precipitated with cold anhydrous ether. After centrifugation, the supernatant was discarded, and the pellet was washed with chilled ether and air-dried. The white precipitate was dissolved in ddH_2_O and purified by preparative reversed-phase high-performance liquid chromatography (RP-HPLC) using an Agilent 1260 Series system. The RP-HPLC was eluted with 0−20% methanol/water with 0.1% TFA at a flow rate of 4 mL min^-1^, and the absorbance of eluates was monitored at 214 nm.

### Standard calibration curve

A standard curve was made using coenzyme A sodium salt hydrate (CoA). The CoA at two-fold serial dilution from 10 to 0 µM (4 µL) was treated with the reaction mixture under the following conditions in a final well volume of 40 µL: 25 mM HEPES (pH 7.5), 150 mM NaCl, 0.01% Triton, and 15 µM ThioGlo4. The fluorescence intensity was measured on a BMG CLARIOstar microplate reader with an excitation wavelength of 400−415 nm, an emission wavelength of 460−485 nm, gain value of 900, the focal height of 6.2 mm using endpoint mode. The baseline was subtracted, arbitrary fluorescence units (AFU) were plotted against the concentration of CoA, an equation of linear regression (y = 5176x + 344.6) was generated, where Y is the AFU and X is the concentration of CoA. All experiments were performed in triplicate.

### Continuous fluorescence-based acetylation assay

A fluorescence assay was adapted to study the kinetics of NatD, which monitors the formation of a ThioGlo4-thiol adduct that exhibits a strong fluorescence at 465 nm.^18,19^ The optimal NatD concentration was investigated by determining the initial rates at different concentrations of the enzyme (25−200 nM) at constant concentrations of H4-8 (50 μM) and AcCoA (50 μM). The reaction was performed at 25 or 37 °C. NatD activity was measured under the following conditions in a final well volume of 40 µL: 25 mM HEPES (pH 7.5), 150 mM NaCl, 0.01% Triton, 25−200 nM NatD, 15 µM ThioGlo4, and 50 µM AcCoA. ThioGlo4 was prepared as a 10 mM stock solution in DMSO, stored, and protected from light at −20 °C. Components except for peptide substrate were initially mixed in a 384-well plate, and the reaction mixture (36 µL) was incubated at 25 or 37 °C for 10 min. The reaction was initiated with the addition of 50 μM H4-8 (4 µL). Fluorescence was monitored on a BMG CLARIOstar microplate reader with excitation of 400−415 nm and emission of 460−485 nm. The standard curve (y = 5176x + 344.6) was used to convert AFU to the concentration of CoA. All experiments were performed in duplicate.

### K_m_ Determination for Peptide Substrate and AcCoA

NatD activity was monitored as one substrate was fixed at different concentrations of the second substrate. To determine K_m_of H4-8, the H4-8 concentration was varied from 0 to 25 μM at fixed AcCoA concentrations (50 μM). We dispensed 36 μL of enzyme mixture (NatD, AcCoA, ThioGLo4 at final concentrations of 0.1 μM, 50 μM, and 15 μM, respectively) into a 384-well plate and incubated at 25 °C for 10 min. To initiate the reaction, 4 μL of varying H4-8 (two diluted from 25 μM to 0.1 μM) solution was added to all wells for a final well volume of 40 µL. Fluorescence was monitored on a BMG CLARIOstar or Neo2 microplate reader (BioTek) with an excitation of 400–420 nm and emission of 465–485 nm for 10 min. The data for the initial rates were fit to the Michaelis-Menten or Substrate inhibition model using least-squares nonlinear regression with GraphPad Prism 7 software. All experiments were performed in duplicate.

### DMSO effect Determination

The effect of DMSO on NatD activity was performed under the following conditions in a final well volume of 100 μL: 25 mM HEPES (pH 7.5), 150 mM NaCl, 0.01% Triton, 0.1 µM NatD, 15 µM ThioGlo4, and 5 µM AcCoA, 4 μM H4-8. The DMSO (1–8 µL) was added to the reaction mixture containing 25 mM HEPES (pH 7.5), 150 mM NaCl, 0.01% Triton, 0.1 µM NatD, 15 µM ThioGlo4, and 5 µM AcCoA. After 10 min incubation, the reactions were initiated by the addition of 10 μL of 40 μM H4-8. Fluorescence was monitored on a BMG CLARIOstar microplate reader with excitation at 400 nm and emission at 465 nm. Data were analyzed using GraphPad Prism 7 software. All experiments were performed in duplicate.

### Stability Evaluation of Protein and Reagent

The stability of the protein and other reagents was assessed by incubating the protein with other components (described below) for 0.5, 1, and 2 hours. Two columns of a NUNC 384-well polystyrene microplate were used. One column served as the negative control (with NatD enzyme added), and another column served as positive controls (no enzyme added). The assay was conducted under the following conditions in a final well volume of 20 μL: 25 mM HEPES (pH 7.5), 150 mM NaCl, 0.01% Triton, 0.1 µM NatD, 15 µM ThioGlo4, and 10 µM AcCoA, 4 μM H4-8. For negative control, 10 μL of premixture that contains 0.2 μM NatD, 30 μM ThioGlo4 and 20 μM AcCoA in the assay buffer (25 mM HEPES pH 7.5, 150 mM NaCl, 0.01% Triton X-100) was dispensed to each well of one column of the plate using Multidrop (Thermo Fisher). For positive control, 10 μL of premixture containing 30 μM ThioGlo4 and 20 μM AcCoA in the assay buffer was dispensed to each well of another column. After 0.5, 1, or 2-hour incubation, the reactions were initiated by the addition of 10 μL of 8 μM H4-8 in the assay buffer. Fluorescence was monitored on a Neo2 microplate reader (BioTek) with an excitation of 400–420 nm and emission of 465–485 nm for 10 min with an interval of 1 min 15 sec at 25 °C. Data were processed in Gen5 software.

### Assay Performance

For assay performance in full 384-well plates, the premixtures of negative and positive controls as described above were dispensed to each well of two 384-well plates, respectively. After 10 min incubation, the reaction was initiated by 10 μL of 8 μM H4-8. The Z’-factor was calculated using equation (1),^20^ where σ _p_ and σ _n_ are the standard deviation of fluorescence signal values of the positive and negative controls, respectively; and µ_p_ and µ_n_ are the mean slope of fluorescence signal values of the positive and negative controls, respectively.

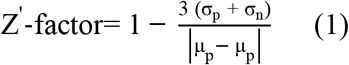

### High throughput screening

Library plates that contain compounds (10 or 1 mM in DMSO) were stored at −80 °C. For plate design, columns 1–2 serve as negative controls (no compound, with NatD enzyme added), and columns 23–24 act as positive controls (no compound, no enzyme added). The test compounds were located in columns 3–22. Library compounds (200 nL for 1 mM compounds or 20 nL for 10 mM compounds) were transferred into columns 3–22 of NUNC 384 well polystyrene microplates using an Echo 555 acoustic liquid dispenser. A total volume of 10 μL of premixture that contains 0.2 μM NatD, 30 μM ThioGlo4 and 20 μM AcCoA in the assay buffer (25 mM HEPES pH 7.5, 150 mM NaCl, 0.01% Triton X-100) was dispensed to each well of columns 1–22 of NUNC plates using Multidrop (Thermo Fisher). Columns 23 and 24 served as positive controls was added 10 μL of premixture containing 30 μM ThioGlo4 and 20 μM AcCoA in the assay buffer. The plates were incubated at room temperature for 10 min. Then, 10 μL of 8 μM H4-8 in the assay buffer was dispensed to initiate the reaction. The plates were quickly spun at 500 rpm for 1 min to ensure the solution sinks to the bottom of microplate wells. Fluorescence was monitored on a Neo2 microplate reader (BioTek) with an excitation of 400–420 nm and emission of 465–485 nm for 10 min with an interval of 1 min 15 sec at 25 °C. Data were processed in Gen5 software.

### MALDI-MS Acetylation and Inhibition Assay

MALDI-MS acetylation and inhibition assay were performed using a Sciex 4800 MALDI TOF/TOF MS in a final well volume of 20 µL: 0.1 µM NatD, 10 µM AcCoA, 4 µM H4-21 peptide, and 10 µM compounds in 25 mM HEPES (pH 7.5) and 150 mM NaCl. Compounds showing >40% inhibition were cherry-picked into column 3 of NUNC 384-well polystyrene microplate using an Echo 555 acoustic liquid dispenser. 10 μL of positive control premixture containing 20 μM AcCoA in 25 mM HEPES (pH 7.5) and 150 mM NaCl was dispensed to each well of columns 4–5 of the plate. 10 μL of enzyme premixture containing 0.2 μM NatD and 20 μM AcCoA in 25 mM HEPES (pH 7.5) and 150 mM NaCl was dispensed to each well of columns 1–3 of the plate. The reaction mixture was incubated at room temperature for 10 min. To initiate the reaction, 10 μL of 8 μM H4-21 in 25 mM HEPES (pH 7.5), and 150 mM NaCl was dispensed to each well of columns 1–5 of the plate. The plates were briefly spun at 500 rpm for 1 second to ensure the solution sinks in the well. After 10 min incubation at room temperature, the samples were quenched in a 1:1 ratio with a quenching solution (10 mg of 2,5-dihydroxybenzoic acid in 1 mL 30%MeCN/70%H_2_O/0.1%TFA (v/v/v)).

The inhibition assay was performed in a final well volume of 100 µL: 0.1 µM NatD, 10 µM AcCoA, 4 µM H4-21 peptide, and varying concentrations of **P1401K09** in 25 mM HEPES (pH 7.5) and 150 mM NaCl at room temperature for 10 min before the addition of 4 µM H4-21 peptide to initiate the reaction. After 10 min incubation at room temperature, the samples were quenched in a 1:1 ratio with a quenching solution (10 mg of 2,5-dihydroxybenzoic acid in 1 mL 30%MeCN/70%H2O/0.1%TFA (v/v/v)). Samples were analyzed by MALDI-MS. Data were processed in Data Explorer.

### IC_50_ determination

NatD activity was measured under the following conditions in a final well volume of 100 µL: 25 mM HEPES (pH 7.5), 150 mM NaCl, 0.01% Triton, 0.1 µM NatD, 15 µM ThioGlo4, and 10 µM AcCoA.^18,19^ The inhibitors in DMSO were 3-fold serially diluted from 300 µM to 0.03 µM. After 10 min incubation at 37 °C, reactions were initiated by adding 10 µM H4-8. All the IC_50_ values were determined in triplicate. Fluorescence was monitored on a BMG ClarioStar microplate reader with excitation 400–415 nm and emission 460–485 nm. Data were processed by using GraphPad Prism software 7.0.

## Results and Discussion

### NatD Protein Purification

Recombinant His-SUMO-NatD was expressed in E. coli overnight and purified by immobilized metal ion affinity chromatography. The His-SUMO tag was removed with SUMO protease to yield untagged NatD in a yield of 12 mg/L of cell culture after purification.

### Fluorescence-based NatD activity assay

This assay monitors the production of CoA in the acetylation reaction by forming a CoA-ThioGlo4 adduct that emits a strong fluorescence at 465 nm (Figure 1A and 1B). We chose ThioGlo4 as the fluorescent probe because it has real-time thiol detection properties, including fast reaction rate, nanomolar sensitivity, and a large signal-to-noise ratio compared to other reported fluorescent probes.^21,22^ Before the kinetic characterization of NatD, we optimized the previously published buffer condition of 25 mM HEPES pH 7.0, 100 mM NaCl, and 1 mM Dithiothreitol (DTT) for our fluorescence assay.^9^ We removed the reducing agent DTT since it can compete with CoA to interact with ThioGlo4. The concentration of NaCl and the pH were optimized to 150 mM and 7.6, respectively, as in the physiological conditions. We compared the optimized buffer condition of 25 mM HEPES pH 7.6, 150 mM NaCl in the presence or absence of detergent (Figure 2A and 2B). The buffer containing detergent displayed a 10-fold increase of fluorescent signal for positive control and a more stable signal for negative control than the buffer without detergent. Thus, the buffer containing detergent was used for NatD characterization. The effect of varying concentrations of ThioGlo4 on NatD was determined to ensure ThioGlo4 does not have any inhibitory activity against NatD. As shown in Figure 2C, ThioGlo4 at 15 uM shows a similar increase of fluorescence intensity compared to that of 5 uM, indicating that ThioGlo4 does not have inhibitory activity against NatD. To ensure the product CoA can be detected in a fast reaction rate, the highest concentration of ThioGlo4 (15 μM) was applied for the fluorescence assay. To design suitable assay conditions for characterizing NatD inhibitors, we determined the steady-state kinetic parameters of both substrates for NatD using a continuous fluorescence assay.^18,19^ Histone H4 peptide substrate was designed based on the first eight amino acids from N-terminus of H4 protein, which was synthesized using standard Fmoc chemistry. For the fluorescence assay, the concentration of CoA formed during the acetylation reaction was derived from a standard calibration curve generated with CoA and ThioGlo4 (Figure 2D). The linearity of the acetylation reaction for NatD concentration was examined in the range of 25 to 200 nM (Figure 2E). We chose 100 nM NatD to carry our kinetic studies in a 384-well plate of 40 µL reaction mixture containing 25 mM HEPES buffer (pH 7.5), 150 mM NaCl, 0.01% Triton X-100, 15 µM ThioGlo4 at 37 °C on BMG CLARIOstar. The linearity of the acetylation reaction to time is shown in Figure 2F. Since room temperature is more amenable for HTS, we retested NatD at concentrations (25 to 200 nM) at room temperature to confirm the optimized condition for HTS. Our results exhibited a linear relationship between NatD concentration and CoA formation at room temperature. The K_m_ value of AcCoA and peptide substrate H4-8 were determined to be 0.5 ± 0.06 µM and 4.0 ± 0.7 µM for NatD, respectively (Figure 2G and 2H). Then we investigated the effects of DMSO on NatD activity by varying the percentage of DMSO from 1–8%, as all small-molecule compounds were stored in DMSO. As shown in figure 2I, NatD exhibited similar enzymatic activities in the presence of 1–8% DMSO, suggesting that NatD is insensitive to DMSO’s effect under the screening conditions with 0.1% of DMSO.

**Figure 1.**
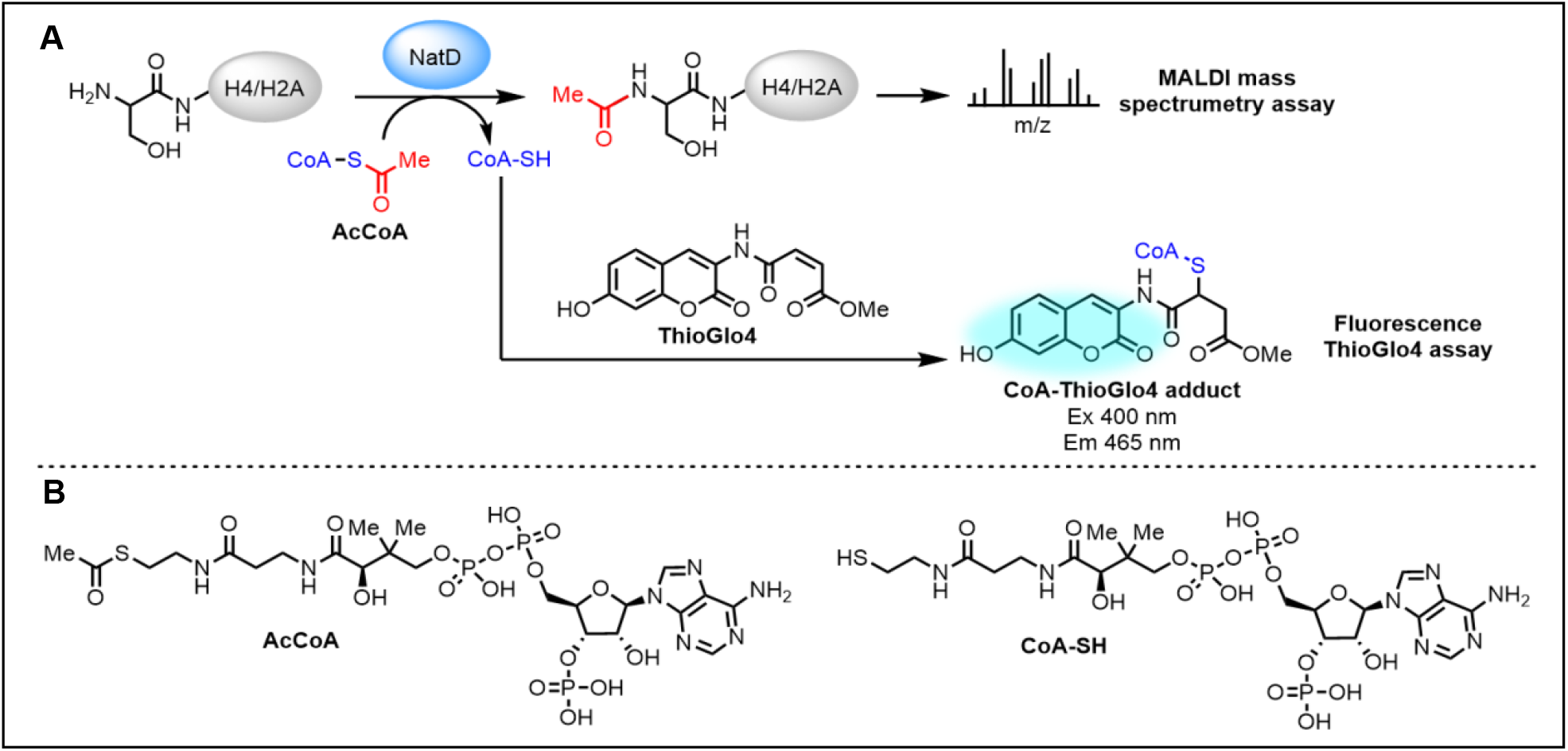
(A) Primary fluorescence ThioGlo4 and MALDI mass spectrometry assays. (B) Selected structures of assay components.

**Figure 2.**
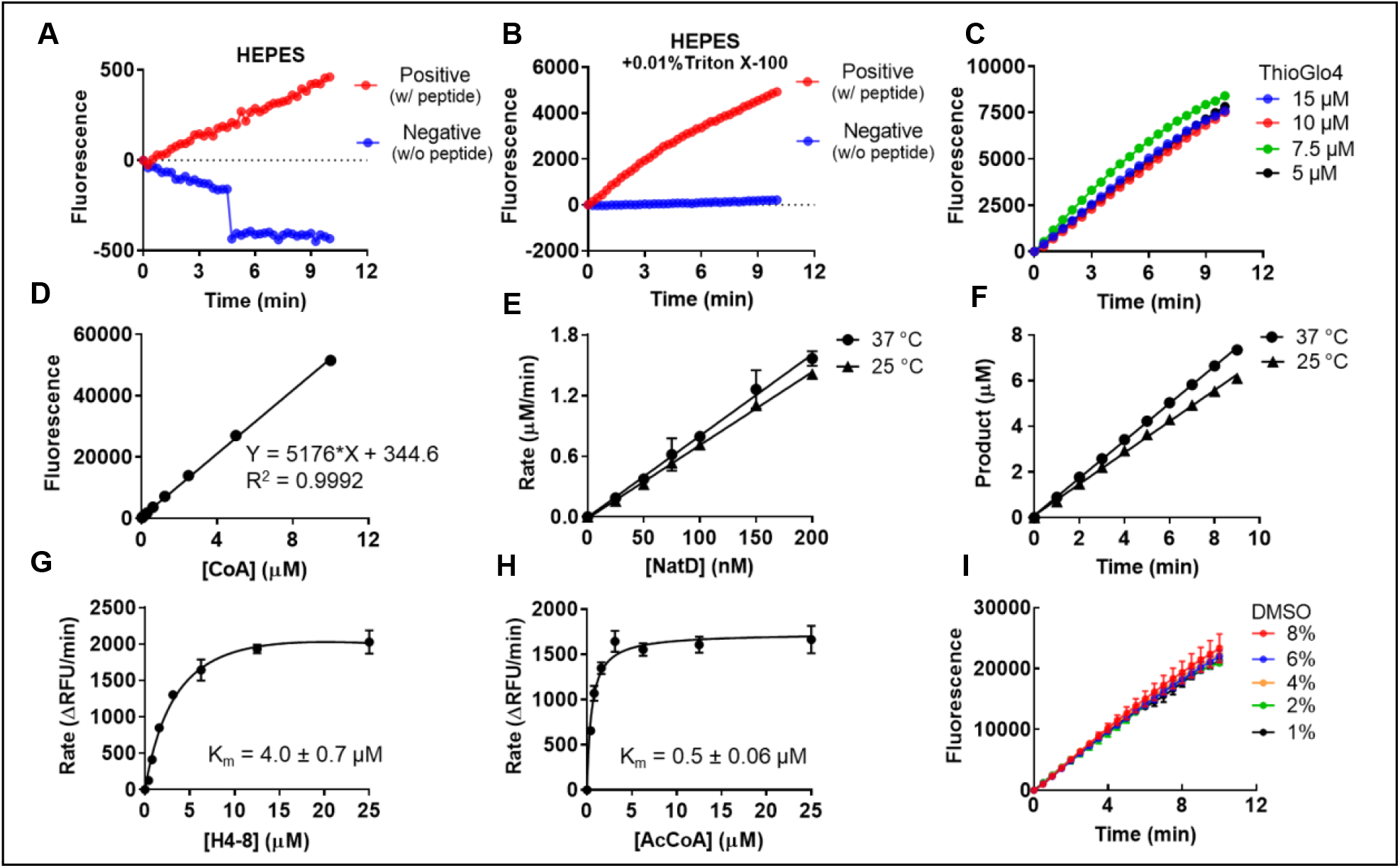
Fluorescence-based NatD activity assay. (A) (B) The effect of buffer (HEPES with or without 0.01% Triton X-100 in NatD activity. (C) The effect of Thioglo4 (5, 7.5, 10, and 15 μM) on NatD activity. (E) Linear relationship between NatD concentration and initial velocity at 37 °C (y = 8.05x − 0.003739, *R*^2^ = 0.997) and 25 °C (y = 7.234x − 0.01427, *R*^2^ = 0.9984). (F) The linear increase in the concentration of products with time at 37 °C (y = 0.8172x + 0.09484, *R*^2^ = 0.9995) and 25 °C (y = 0.6855x + 0.09642, *R*^2^ = 0.9981). (I) The effect of % of DMSO on NatD activity.

### Optimization for Primary Fluorescence Assay in a 384-Well Format

Based on the results from BMG CLARIOstar, we further miniatured the assay from a total volume of 40 µL to 20 µL in a 384-well plate on a BioTek Synergy Neo2 microplate reader. We re-determined the K_m_ values of both H4-8 peptide and AcCoA under modified condition, yielding 3.5 ± 0.7 µM and 1.1 ± 0.2 µM, respectively (Figure 3A and 3B). Comparable K_m_ values for both H4-8 and AcCoA confirmed the reproducibility despite data obtained from two different instruments. Since the NATs utilize the AcCoA as their acetyl donor, AcCoA competitive inhibitors may impose a challenge for NATs’ selectivity. Because selectivity of any given inhibitor is essential to probe its respective target functions, our assay conditions are designed to identify selective inhibitors for NatD by targeting its unique peptide substrate binding site.^9^ Therefore, the screening was performed using H4-8 at a K_m_ value concentration (4 μM) and AcCoA at a saturating concentration (10 μM) to favor the condition to identify compounds targeting peptide-binding site instead of AcCoA binding site.

**Figure 3.**
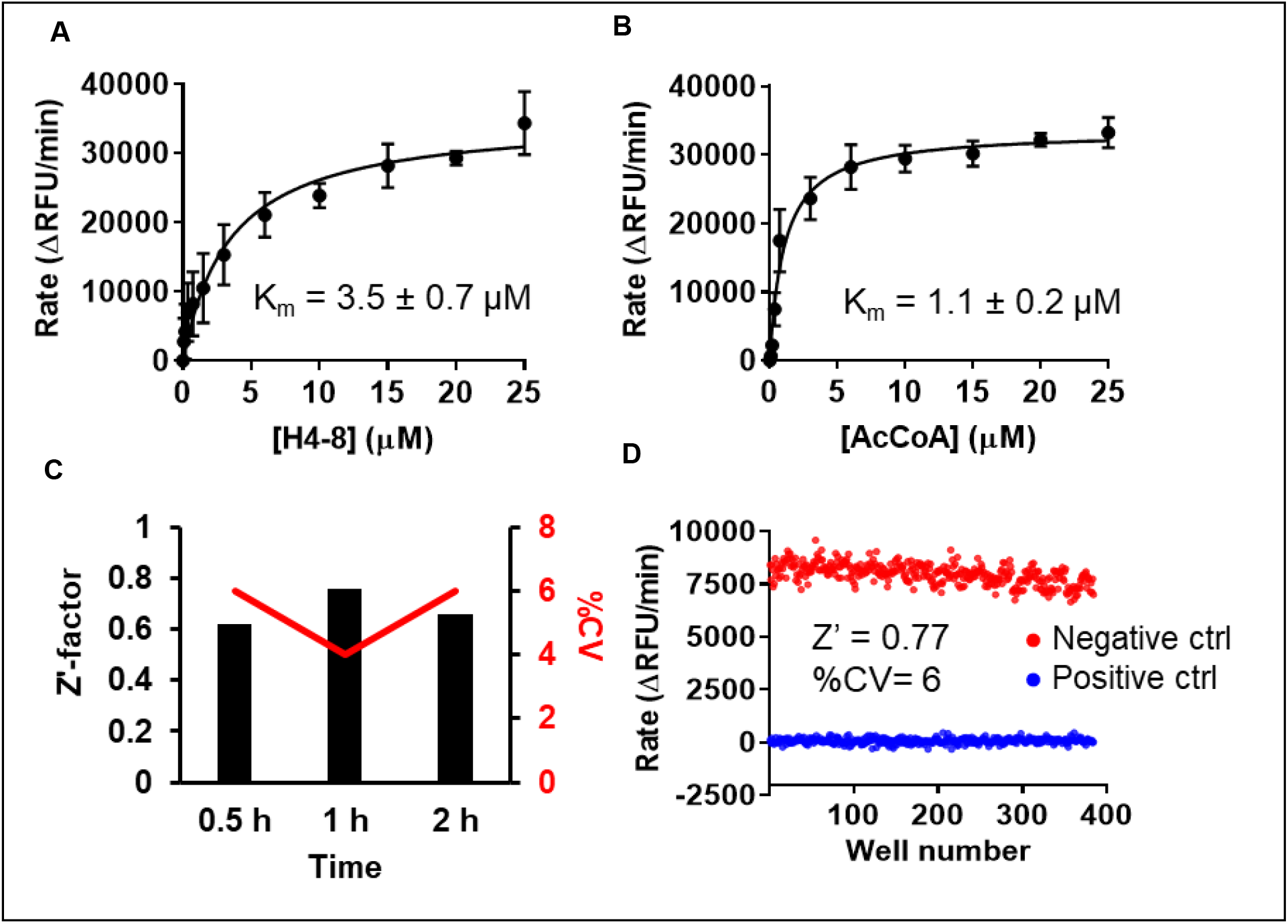
Optimization for fluorescence ThioGlo4 assay. (A)(B) Re-determination of Km values of H4-8 and AcCoA in 384-well format using BioTek Synergy Neo2 microplate reader. (C) Assay performance (Z’-factor and %CV) at incubation time of 0.5 h, 1 h, and 2 h. (D) Assay performance in a 384-well format. Red dots: 384 wells of negative controls. Blue dots: 384 wells of positive controls.

### Stability of Protein and Reagent

To check the reagent and protein stability, we investigated the assay performance of reaction mixtures preincubated for 0.5, 1, and 2 hours under the optimized condition (0.1 μM of NatD, 4 µM of the H4-8, 20 µM of AcCoA, and 15 µM of ThioGlo4). Sample wells containing 0.1 μM NatD served as negative controls, while those wells without NatD served as positive controls for inhibition. This assay displayed a robust performance with an average Z’-factor of 0.68 and CV of 5 % after 0.5, 1, or 2-hour preincubation (Figure 3C).

### Assay Performance

To evaluate the assay’s robustness as an HTS-feasible assay, we performed two full 384-well plates of one positive control plate and one negative control plate under the optimized condition using automated liquid handlers (Multidrop) for dispensing. A Z’-factor of 0.77 and a CV of 6% were obtained (Figure 3D).

### Pilot screen with LOPAC library

We then applied the optimized condition to perform the pilot screen with the LOPAC 1280 library in order to investigate its applicability in HTS (Figure 4A). The compounds were screened at a final concentration of 10 µM in 0.1% DMSO under the optimized condition. Initial hits were defined as those active compounds that displayed over 40% inhibition. Among 1280 compounds, twelve compounds met our hit criteria in the pilot screen (Figure 4B). Meanwhile, we examined day-to-day and plate-to-plate variability by placing two columns (32 samples) of negative and positive controls in each plate as internal controls. These internal controls exhibited an average Z’-factor of 0.82 and CV of 5%, validating the reproducibility of this fluorescence-based assay for HTS (Figure 4C).

**Figure 4.**
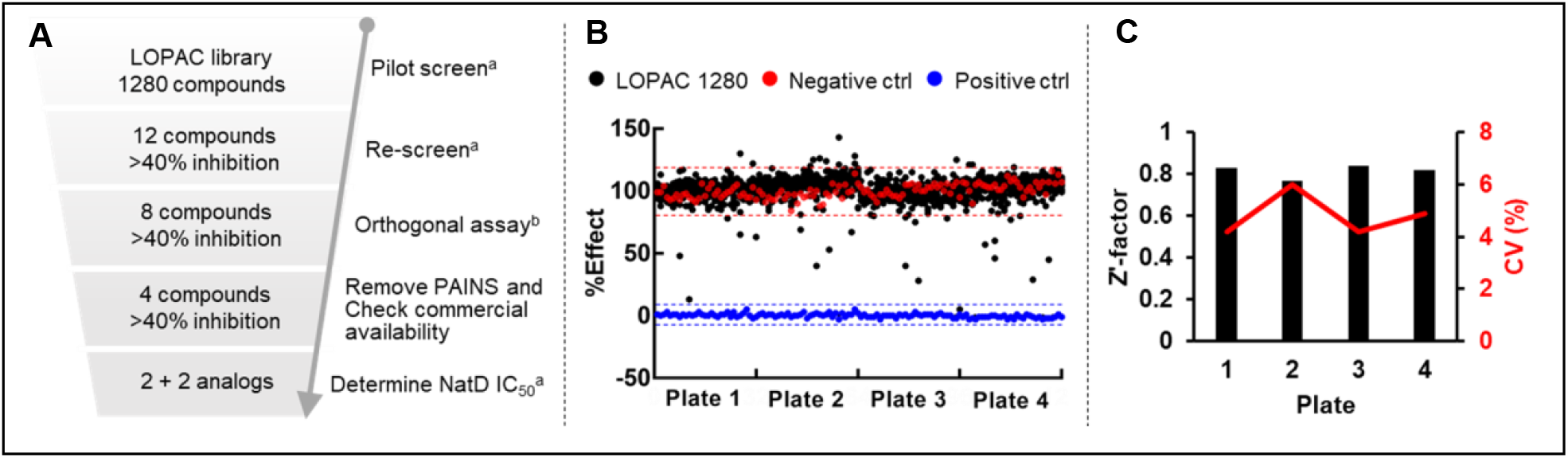
Pilot screen summary in the LOPAC library. (A) HTS screening flowchart. ^a^Performed by fluorescence assay. ^b^Performed by MALDI-MS assay (B) Screening results. Red and blue dash lines represent 3 standard deviations from the mean of negative and positive controls, respectively. (C) Z’-factors and %CV of each plate in the pilot screen.

### Validation Assays

To confirm the initial hits identified in the primary screen, those cherry-picked twelve compounds were retested their inhibition at 10 µM in the fluorescence assay. Among twelve screened compounds, eight compounds reproducibly exhibited >40% inhibition (Table 1). Next, we conducted an orthogonal MALDI-MS assay to directly monitor acetylated peptide formation to remove false positive hits that interfere with the fluorescent dye.^23^ H4-21 peptide was chosen because it showed higher signal intensity than H4-8 in the MALDI-MS assay. Triton X-100 was removed from the reaction buffer to prevent it from suppressing the MALID-MS signal.^24^ Of the eight compounds that exhibited repeatable activity in the fluorescence assay, four displayed >40% inhibition in MALDI-MS assay are aurintricarboxylic acid (ATA, **P1401K09**), tyrphostin B44(-) (**P1404L15**), reactive blue 2 (**P1404C10**), and cisplatin (**P1403H18**) (Table 1).

**Table 1.**
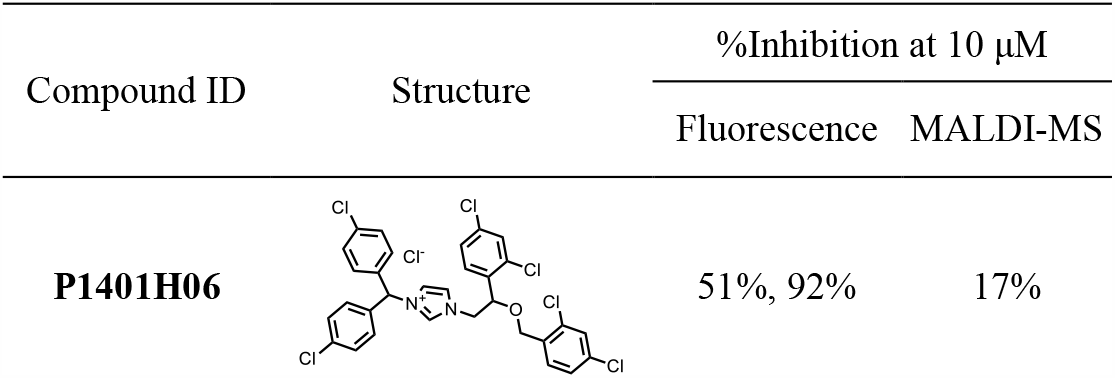
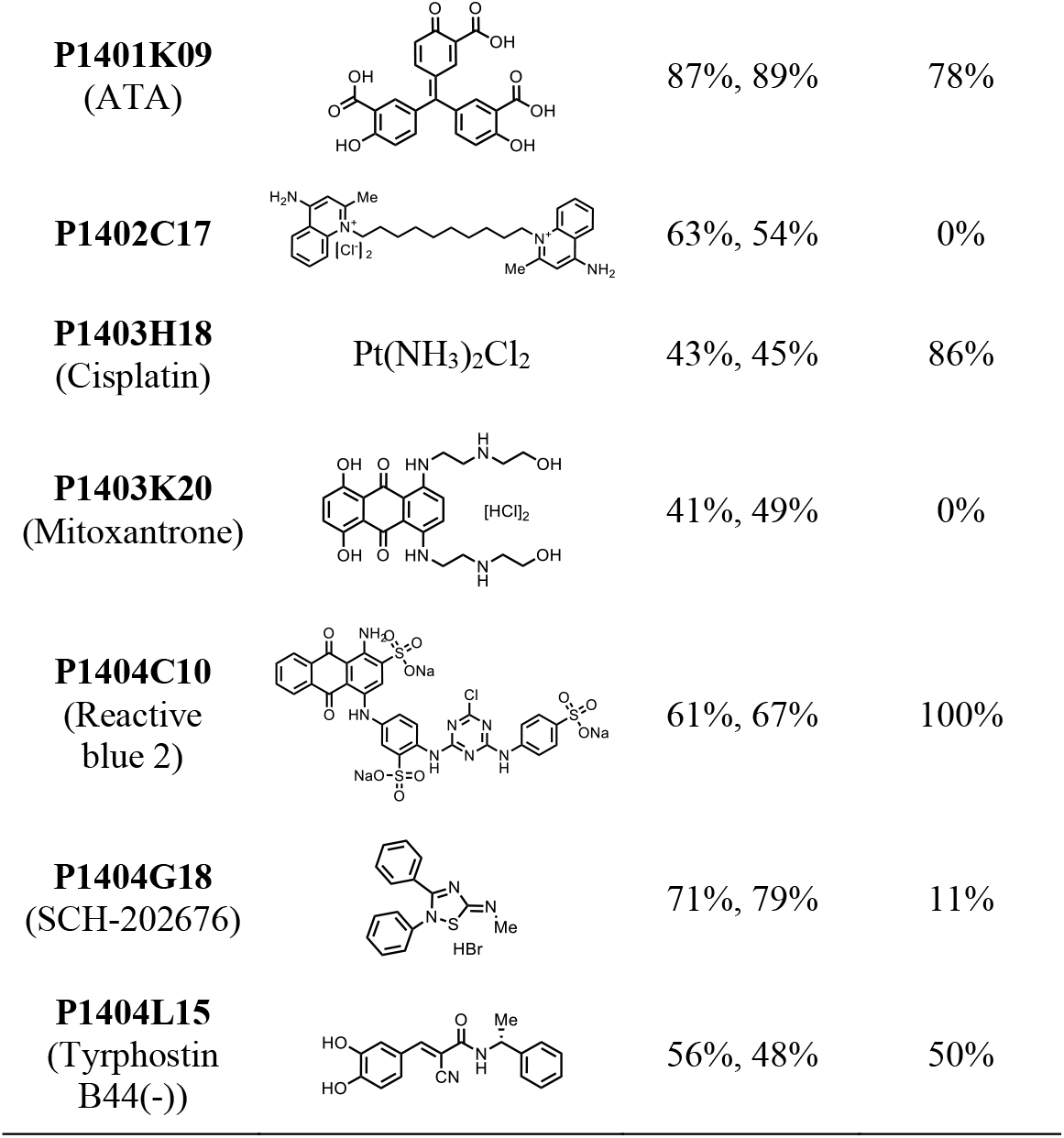
%Inhibition at 10 μM of hits from LOPAC pilot screen.

| Compound ID | Structure | %Inhibition at 10 μM | |
| --- | --- | --- | --- |
|  |  | Fluorescence | MALDI-MS |
| **P1401H06** |  | 51%, 92% | 17% |
| **P1401K09**<br>(ATA) |  | 87%, 89% | 78% |
| **P1402C17** |  | 63%, 54% | 0% |
| **P1403H18**<br>(Cisplatin) | Pt(NH<sub>3</sub>)<sub>2</sub>Cl<sub>2</sub> | 43%, 45% | 86% |
| **P1403K20**<br>(Mitoxantrone) |  | 41%, 49% | 0% |
| **P1404C10**<br>(Reactive blue 2) |  | 61%, 67% | 100% |
| **P1404G18**<br>(SCH-202676) |  | 71%, 79% | 11% |
| **P1404L15**<br>(Tyrphostin B44(-)) |  | 56%, 48% | 50% |

### IC_50_ characterization

Although ATA is known as a promiscuous inhibitor for multiple targets and considered a Pan-Assay Interference (PAINS) compound,^25^ we selected ATA for further characterization as it can be used as a positive control to assess assay performance. We were able to source ATA, tyrphostin B44(-), and its analogs (tyrphostin AG 835 and AG 490) to characterize their IC_50_ values (Figure 5). ATA showed inhibitory activity against NatD at 1.5 ± 0.1 µM in the fluorescence assay and 2.9 µM in the MALDI-MS assay (Figure 6). Tyrphostin B44(-) showed IC_50_ of 63 ± 4 µM against NatD. AG 835, a diastereomer of Tyrphostin B44(-), displayed a 2-fold increased inhibition (IC_50_ = 38 ± 2 µM) against NatD, suggesting the chirality of the methyl group affects NatD inhibition. AG 490 without a methyl group on α,β-unsaturated ketone showed a comparable NatD inhibition with Tyrphostin B44(-).

**Figure 5.**
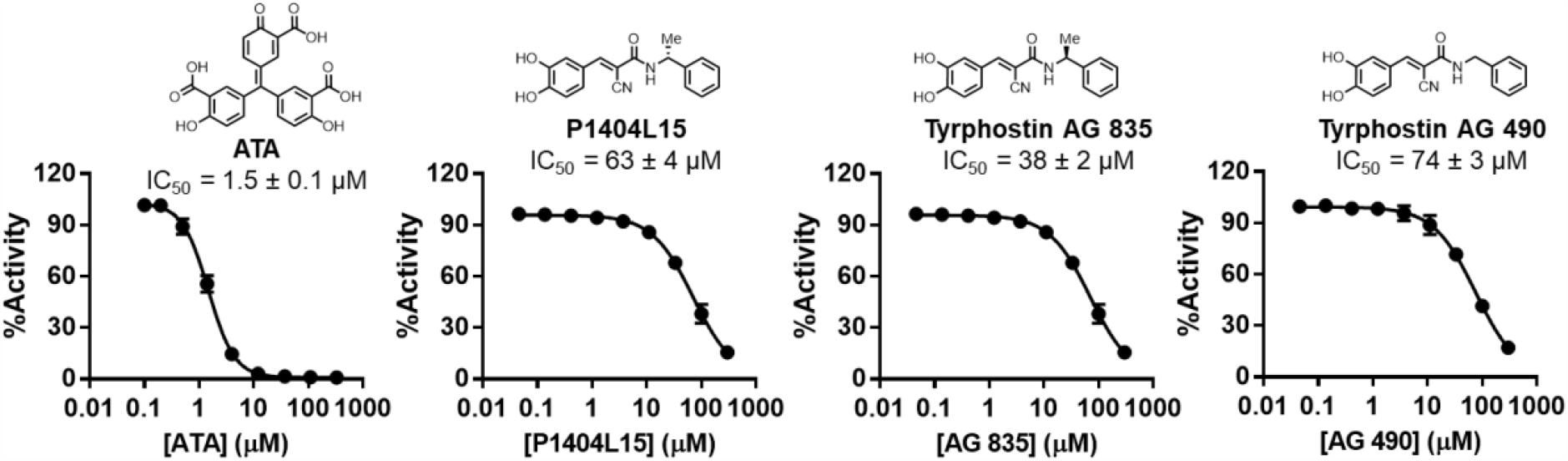
Fluorescence IC_50_ characterization of selected hits and analogs from LOPAC pilot screen (n = 2).

**Figure 6.**
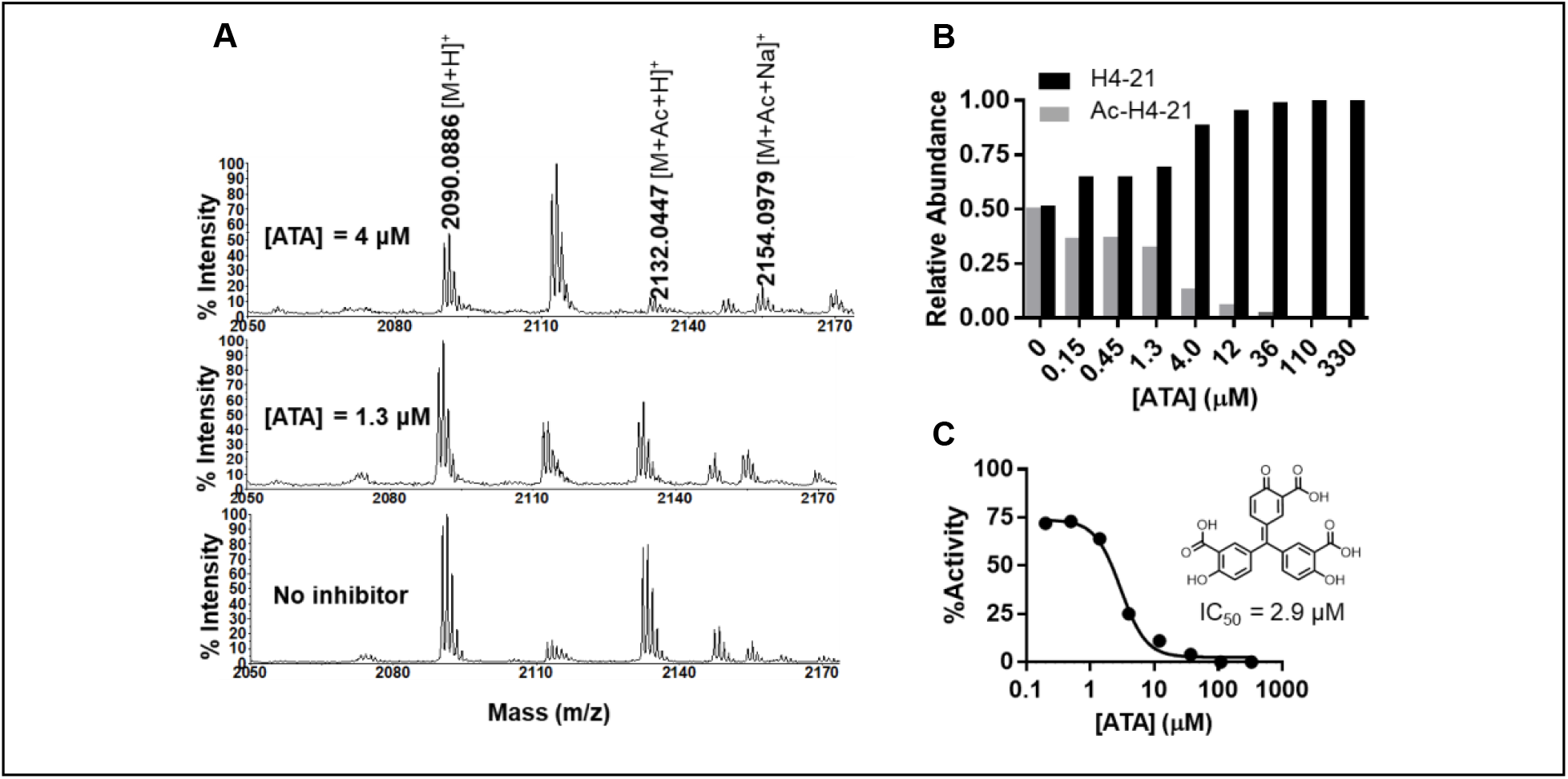
MALDI-MS acetylation inhibition assay for **ATA** (n=1). (A) Representative MS at 0, 1.3, and 4 µM of ATA. (B) Quantification of ATA inhibition. (C) IC_50_ of ATA.

## Conclusion

In this study, we established and miniaturized a continuous fluorescence-based assay to monitor the activity of NatD in a 384-well format. This assay showed favorable Z’-factors, which enabled us to conduct a pilot screen of the LOPAC library and identify small molecule inhibitors. The pilot screen showed high reproducibility and Z’-factors > 0.77, demonstrating this assay’s robustness for HTS. In the fluorescence-based assay, ThioGlo4 is used to react with the CoA to form a thiol adduct, generating a fluorescence signal. The fluorescent dye may interfere with autofluorescence compounds or react with screened compounds to yield false-positive signals. Therefore, we perform a MALDI-MS counter assay to overcome this problem, directly monitoring the acetylated peptide without using fluorescence dye. Among two confirmed hits, ATA can be used as a positive control for monitoring the reproducibility of the assay performance. **P1404L15** and its two analogs showed modest inhibition, which will be examined further for optimization. In conclusion, we demonstrated the feasibility of the continuous fluorescence-based assay for HTS to identify NatD small molecule inhibitors, which could facilitate the future discovery of NatD inhibitors to investigate the biological functions of NatD. Notably, this assay has the potential to be applied for characterization of other NATs.

## Acknowledgments

We thank the Chemical Genomics Facility in the Institute for Drug Discovery at Purdue University for the compound library and robotic instruments.

